# PIWI proximity proteome reveals Set1-mediated piRNA biogenesis for transposon silencing in telomere

**DOI:** 10.64898/2026.03.30.715253

**Authors:** Wakana Isshiki, Hiroko Kozuka-Hata, Masaaki Oyama, Toshie Kai, Taichiro Iki

## Abstract

Silencing complexes formed by PIWI-clade Argonaute (Ago) proteins and PIWI-interacting RNAs (piRNAs) are essential guardians of genome integrity, controlling the deleterious activities of transposable elements (TEs) in animal germline. However, our understanding of PIWI-piRNA-directed TE silencing remains incomplete. Here, we systemically characterize the proximity proteome of PIWI members, Piwi, Aubergine (Aub), and Ago3 in the germline of *Drosophila* ovaries. Functional screening identifies previously uncharacterized factors involved in TE silencing, including H3K4me3 writer and transcriptional coactivator Set1. Transcriptome analysis reveals that Set1 acts as an indispensable repressor for TEs, particularly those forming telomeres. The involvement of Set1 in Piwi pathway is further supported by its critical role in the production of antisense, TE-targeting piRNAs. Notably, catalytic activity of Set1 is dispensable for TE silencing. Genome-wide chromatin binding analysis using CUT&Tag demonstrates that Set1 preferentially associates with TE sequences and localizes to subtelomeric piRNA cluster loci, suggesting a role in promoting piRNA precursor transcription through direct binding. Collectively, these findings uncover a noncanonical function of Set1 in Piwi-mediated TE silencing and telomere control in germline nuclei.

## Introduction

Transposable elements (TEs) are mobile DNA sequences constituting a substantial part of genomes in most organisms (Wells & Feschotte, 2020). Although TEs act as drivers of genomic evolution and species diversification, the harmful potential as mutagens requires the hosts to establish the control mechanisms. The control is particularly important in germline cells where activated TEs accumulate damages on the inheritable genomes and impair the production of functional gametes. In animals, TE control in germline cells relies on silencing mediated by complexes of PIWI clade Argonaute (Ago) family proteins and the sequence-specific guide, PIWI-interacting (pi)RNAs (Wang *et al*, 2023b).

PIWI-piRNA-directed TE silencing has been studied in different animals including the invertebrate model, *Drosophila melanogaster. D. melanogaster* possesses three PIWI members including Piwi, Aubergine (Aub), and Ago3, all of which are expressed in germline cells. Piwi differs from the other two in that it functions in the nucleus. Guided by piRNAs, Piwi complexes recognize the nascent transcripts from target loci, mediating the (co-)transcriptional silencing and heterochromatin formation (Le Thomas *et al*, 2013; Yu *et al*, 2015b; Sienski *et al*, 2015a; Mugat *et al*, 2020; Ariura *et al*, 2024). In contrast, Aub and Ago3 are cytoplasmic proteins, localizing to nuage, a perinuclear membraneless organelle formed in germline cells (Suyama & Kai, 2025; Kawaguchi *et al*, 2025). In nuage, Aub and Ago3 mediate the reciprocal cleavage of targets containing sense and antisense TE sequences. 3’ cleavage products generated by either of the two proteins are passed to the other (i.e., from Aub to Ago3, or from Ago3 to Aub), then processed into a piRNA. Newly generated piRNAs direct the next round of target cleavage and piRNA production. These cycles are called ping-pong, coupling TE degradation and piRNA amplification in germline cells. Because target cleavage occurs between the nucleotides complementary to 10^th^ and 11^th^ nucleotides of guide piRNA, and new piRNA is produced from the 5’ end of cleavage product, germline piRNAs exhibit 10-nt overlap signature (ping-pong signature). In addition to ping-pong, precursor fragments incorporated to Aub can be processed into multiple piRNAs on mitochondrial outer membrane. These phasing/trailer piRNAs are mainly loaded onto Piwi, translocating the complexes to the nucleus (Ge *et al*, 2019). Ping-pong cycles process diverse transcripts including intact TE mRNAs and long non-coding RNAs derived by noncanonical transcription of heterochromatinized, large intergenic regions accumulating truncated TE sequences, defined as piRNA cluster loci (Brennecke *et al*, 2007a).

piRNA-directed TE silencing is achieved through interactions between PIWI proteins and other components. These include nuclear factors involved in Piwi-piRNA-directed target heterochromatinization (e.g., Panx, Nxf2, SetDB1/Eggless) (Zhao *et al*, 2019; Murano *et al*, 2019; Fabry *et al*, 2019; Batki *et al*, 2019; Yu *et al*, 2015b; Sienski *et al*, 2015a). In cytoplasm, Tudor domain proteins (Krimper, Tejas, Tapas, Qin/Kumo) and RNA helicases (Spn-E, Vas) are enriched in nuage and support the ping-pong cycles mediated by Aub and Ago3 (Webster *et al*, 2015; Lin *et al*, 2023; Handler *et al*, 2011; Zhang *et al*, 2011; Anand & Kai, 2012; Qi *et al*, 2011; Saito *et al*, 2010; Lim & Kai, 2007; Kawaguchi *et al*, 2025). Nonetheless, characterizing the interactions between PIWI members and other factors are often challenging because of their dynamics in piRNA biogenesis and TE silencing. Hence, the repertoire of piRNA pathway components is heretofore unelucidated, and our understanding of piRNA-directed TE silencing remains incomplete.

Proximity-dependent biotin labelling (BioID/TurboID) is a powerful tool for characterizing the physical interactions and proximity relationships of proteins *in vivo* (Roux *et al*, 2012; Branon *et al*, 2018; Choi-Rhee *et al*, 2004). These techniques rely on the promiscuous biotin ligase activity exhibited by BirA derived from *Escherichia coli*. TurboID utilizes an efficient and compact BirA variant called mini(m)Turbo established through directed evolution (Branon *et al*, 2018), which we previously introduced in male germline cells to study testis-specific factors (Iki *et al*, 2023; Kai *et al*, 2025; Iki *et al*, 2020). In order to deepen the understanding of piRNA-directed TE silencing, this study characterized the proteins in the physical proximity of individual PIWI proteins in female germline cells where TE silencing has been extensively studied. The obtained proximity proteome of germline PIWI members not only confirmed reported interactions but also revealed the unrecognized links. Moreover, PIWI proximity proteome contained a list of factors which have not been characterized in piRNA pathway thus far. Of those, this study further characterizes Set1, a histone methyltransferase and transcriptional coactivator conserved across eukaryotes. Our data highlight the hitherto unappreciated importance of Set1 in Piwi-piRNA-directed silencing of TEs forming telomeres in *Drosophila* genome.

## Results

### Labelling of PIWI proximity factors and purification from germline cells

To characterize the proximity proteome of PIWI proteins using TurboID, we generated a series of transgenes encoding mTurbo-GFP-fused Piwi, Aub, or Ago3 (Figure 1A). These transgenes were expressed in ovarian germline cells under the control of *nos* promoter activity (UASp/Gal4 binary system using NGT40;nos-Gal4 (NN) driver) (Rørth, 1998; Grieder *et al*, 2000). The biotinylation activities of individual fusion proteins were confirmed by detecting the streptavidin-HRP-dependent signals (Figure S1A). However, pulldown using streptavidin was inefficient and biotinylated proteins largely remained in the flow through, possibly due to interfering factors accumulated during differentiation. To circumvent this technical challenge, we depleted one of the differentiation factors, *bag of marbles* (*bam*) (McKearin & Spradling, 1990), by germline knockdown (GLKD), and collected germline stem cell (GSC)-like cells (GSCLCs) impaired in differentiation (Figure 1A, S1BC). Although TurboID in GSCLCs showed weaker biotinylation signals in the input, the pulldown efficiency was dramatically improved (Figure S1A).

**Figure 1.**
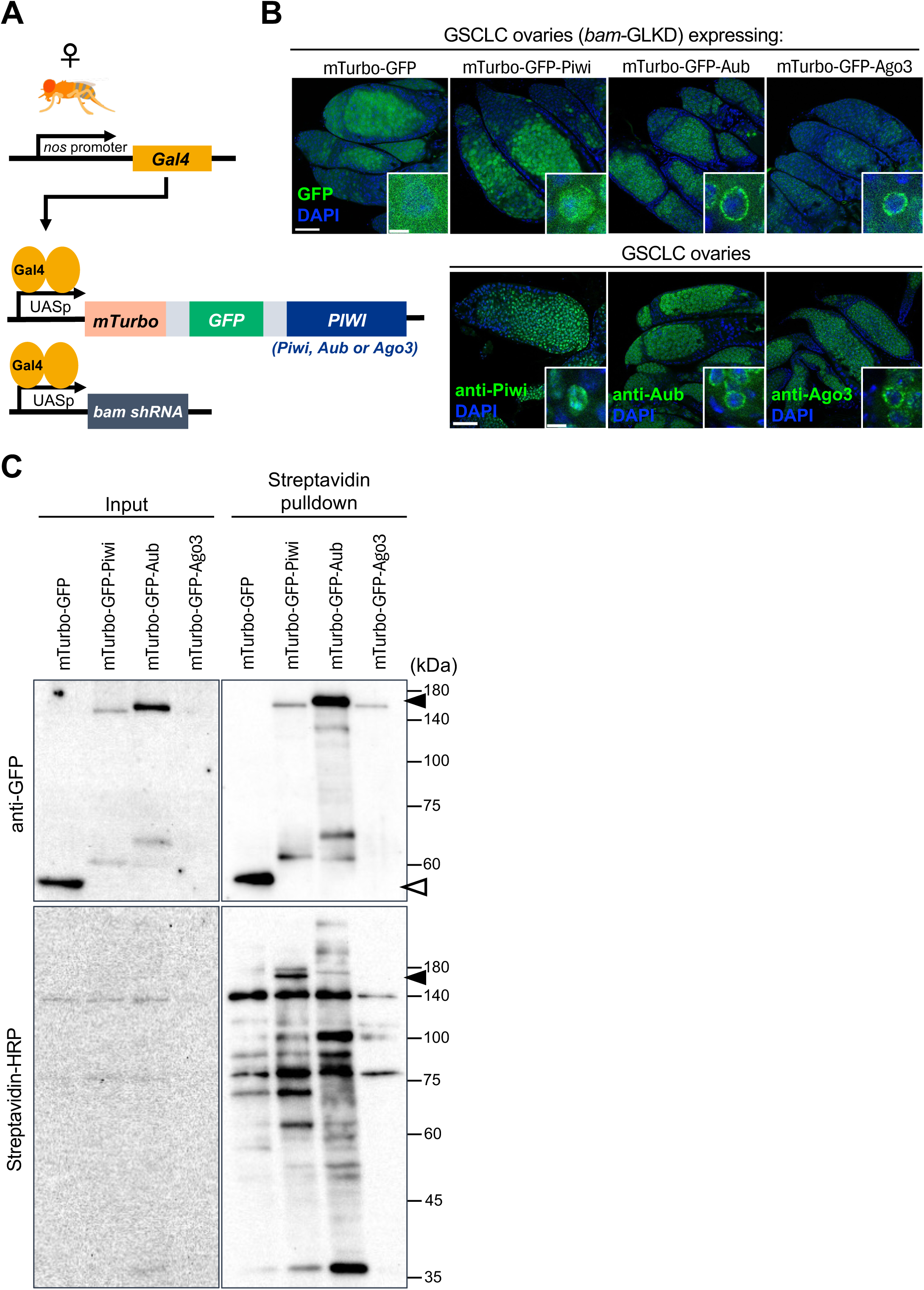
PIWI proximity biotin labelling in germline stem cell-like cells (GSCLCs) (A) Experimental design for PIWI proximity biotin labelling in female germline cells. mTurbo-GFP-Piwi, –Aub, or –Ago3 is expressed in germline cells using UASp/Gal4 system with NGT40; nos-Gal4 (NN) driver. mTurbo-GFP serves as negative control. Depletion of *bam* using short hairpin (sh)RNA enables TurboID in GSCLCs. (B) Subcellular localization of mTurbo-GFP fusion proteins in GSCLC ovary. Upper panels; fluorescent signals from GFP (green). Lower panels; immunostaining signals from endogenous Piwi, Aub, or Ago3 (green). DAPI (blue) for nuclei. Scale bar = 50 μm or 5 μm in the insets. (C) Blotting images for proteins extracted from GSCLC ovaries expressing mTurbo fusion proteins (Input), and the streptavidin-bound fraction (Pulldown). Upper panels; immunoblotting with anti-GFP antibody. Lower panels; blotting with streptavidin-HRP. White and black arrowheads indicate mTurbo-GFP and mTurbo-GFP-PIWI proteins, respectively.

In GSCLCs, mTurbo-GFP alone displayed a broad distribution across both the nucleus and cytoplasm (Figure 1B). In contrast, mTurbo-GFP-PIWI proteins exhibited subcellular localization patterns consistent with those of their endogenous counterparts; Piwi accumulated in the nucleus, while Aub and Ago3 were enriched in the perinuclear nuage. The concordance suggests that N-terminal mTurbo-GFP tagging does not substantially disrupt the interaction networks of individual PIWI proteins. Supporting this notion, N-terminal mTurbo-FLAG fusion preserved Aub functions in male germline (Iki *et al*, 2023). Notably, the steady-state levels of fusion proteins were below those of endogenous counterparts (Figure S1D). mTurbo-GFP-Ago3 was the least stable and barely detectable in the input (Figure 1C, anti-GFP), accounting for the weak streptavidin-HRP signals in Ago3-TurboID condition (Figure 1C). Nevertheless, the streptavidin pulldown displayed distinct signal patterns, suggesting biotinylation-dependent enrichment of specific factors in each PIWI-TurboID condition.

### Characterization of germline PIWI proximity proteomes

Mass spectrometry analysis and the label-free quantification (LFQ) of streptavidin pulldown samples identified the proximity factors of individual PIWI proteins (average abundance ratios >1 compared to the control GFP-TurboID conditions in biological duplicates) (Table S1). These included 37, 33, and 49 factors for Piwi, Aub, and Ago3, respectively (Figure 2A and 2B, S2). Notably, the identified proximity factors contained known piRNA pathway components (Figure 2A, highlighted in yellow, Table S2), and the significant enrichment was confirmed by Gene Ontology (GO) analysis (Figure 2C). These results suggest that the proteome reflects the physical and functional interactions between PIWI members and other factors in germline cells.

**Figure 2.**
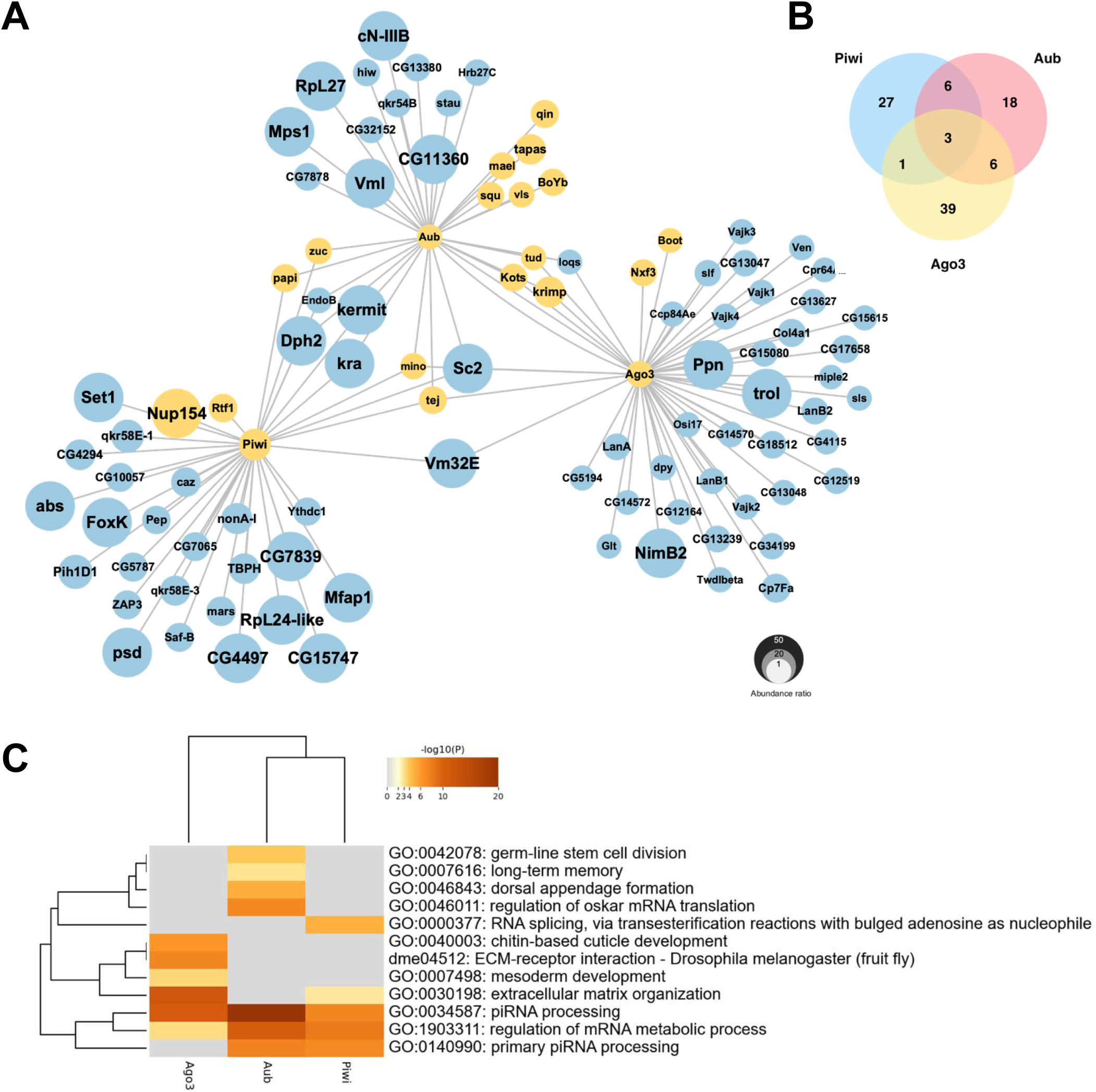
Proximity proteome of PIWI proteins in germline cells. (A) Network diagram summarizing proximity factors of individual PIWI proteins (Piwi, Aub, and Ago3) identified by mass spectrometry (nanoLC-MS/MS) followed by label-free quantification. Yellow nodes: factors known to be involved in piRNA pathway. Size of circle reflects abundance ratio (mTurbo-GFP-PIWI/mTurbo-GFP) based on biological duplicate TurboID data. (B) Venn diagram shows the number of proximity proteins shared by or unique to individual PIWI members. (C) GO enrichment analysis of PIWI proximity factors. Hierarchical clustering and heatmap show significantly enriched GO terms. Color intensity represents the significance of enrichment (-log10 p-value).

### Screening of PIWI proximity factors involved in piRNA biogenesis and TE silencing

Most factors identified in the proximity of PIWI members have not been functionally characterized in piRNA pathway or TE silencing. Hence, we investigated the possible involvement of 54 factors, for which short hairpin-based RNAi lines were available, by germline knockdown (GLKD) screening using NN driver. Given that impaired piRNA biogenesis can be associated with nuage collapse and mislocalization of its components, we examined the localization of Krimp, a key scaffold of nuage (Lim & Kai, 2007; Patil & Kai, 2010). In parallel, we observed the accumulation of Gag protein encoded by a non-LTR retroelement *HeT-A*, as a readout of TE derepression (Shpiz *et al*, 2011). We confirmed that GLKD of *aub* leads to the accumulation of *HeT-A* Gag protein around the oocytes of developing egg chambers (Figure 3A), and the loss of Krimp from perinuclear nuage and the formation of abnormal foci in the cytoplasm (Figure 3B, S3B).

**Figure 3.**
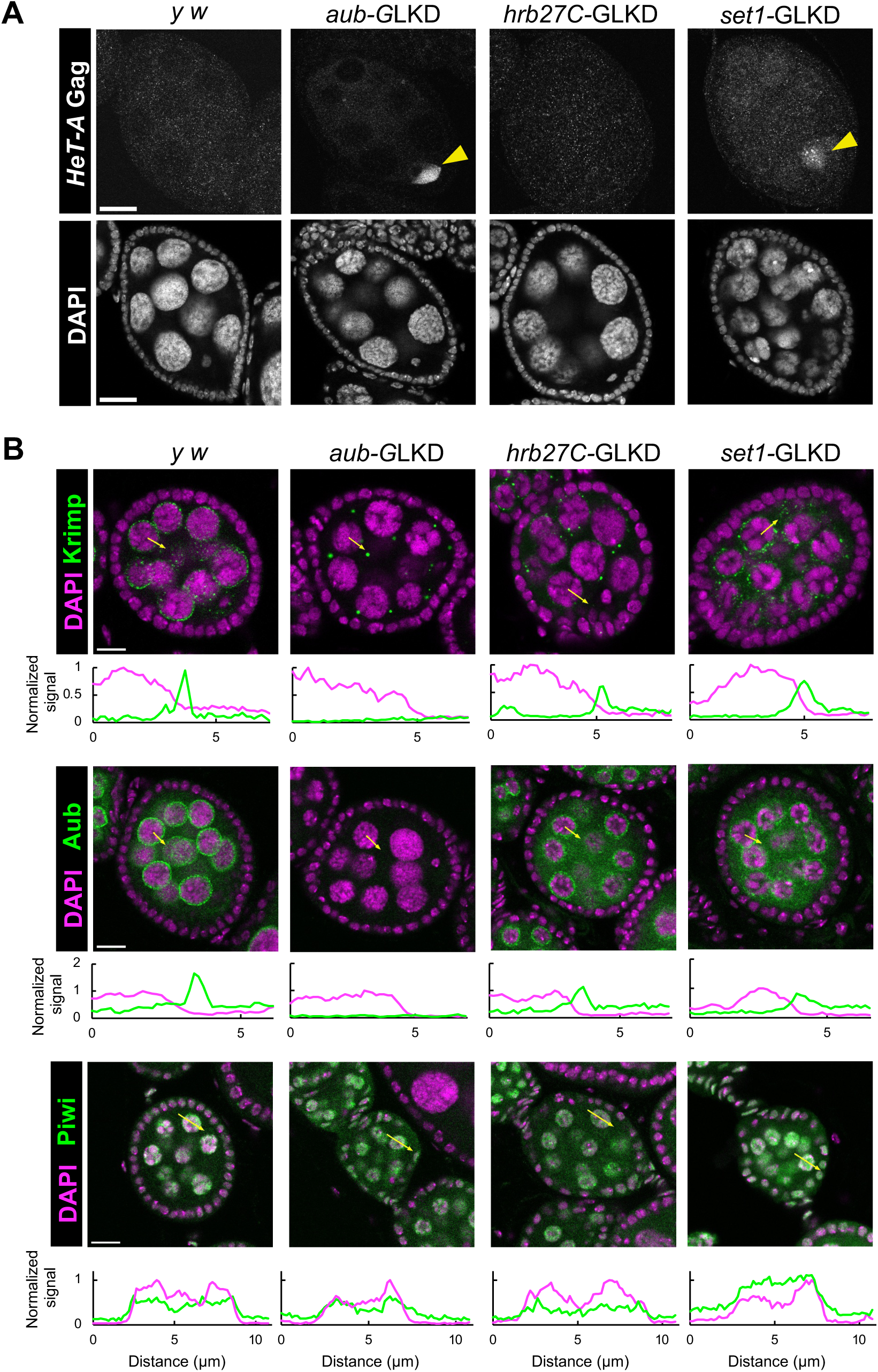
Germline knockdown (GLKD) screening of PIWI proximity factors. (A) Immunofluorescence signals from *HeT-A* Gag protein (top panels) in egg chambers of control (*y w*) or GLKD of indicated genes. DNA stained with DAPI (bottom panels). Arrowheads; Gag proteins accumulating around oocyte. Scale bars = 20 μm. (B) Immunofluorescence signals of Krimp (green) in the egg chambers of control (*y w*) or of indicated GLKD conditions. DNA stained with DAPI (magenta). Fluorescence signals along yellow arrows are plotted in the bottom panels (signals normalized by setting 1 for the highest value of DAPI). Scale bars = 10 μm. (C) Immunofluorescence signals of Aub or Piwi (green) in the egg chambers of indicated conditions.

GLKD screening identified several factors including Hrb27C and Set1, the proximity factor of Aub and Piwi, respectively (Table S3). Both factors are conserved across eukaryotes, and their molecular and physiological functions have been characterized. Hrb27C (also known as Hrp48) is an hnRNP family member required in GSC for the maintenance (Yan *et al*, 2014; Ables *et al*, 2016; Finger *et al*, 2023). Set1 is an enzyme acting inside COMPASS complex responsible for di– and tri-methylation of histone H3 at lysine 4 (H3K4me2/3), a modification associated with transcription start sites (TSSs) for RNA polymerase (pol)II activation (Ardehali *et al*, 2011; Mohan *et al*, 2011; Wang *et al*, 2023a). We observed distinct phenotypes in *hrb27C*-GLKD and *set1*-GLKD. In *hrb27C*-GLKD, abnormal Krimp foci appeared in the cytoplasm, similar to *aub*-GLKD (Figure 3B). However, *HeT-A* Gag protein was under detectable level (Figure 3A). In *set1*-GLKD, though abnormal Krimper foci formation was not clear, *HeT-A* Gag protein accumulated around the oocyte. In both *hrb27c-* and *set1*-GLKD conditions, perinuclear Krimp signals could be weaker but still observed (Figure 3B). Consistently, Aub and Piwi did not show drastic changes in their localization (Figure 3C, S3C). Of note, RT-qPCR confirmed target downregulation in individual GLKD conditions (Figure S3D).

### *set1* and *hrb27C* functionally interact with *piwi* and *aub*, respectively, to control TEs

To investigate the global effect of Hrb27C or Set1 depletion in the germline on TE expression, we compared the transcriptome from *hrb27C*– or *set1*-GLKD ovaries with that from *gfp*-GLKD control ovaries (Figure 4A, S4A). In parallel, *aub*– and *piwi*-GLKD ovary transcriptomes were generated as references for TE derepression. Our analysis included germline stem cell-like cell (GSCLC) condition in which PIWI proximity proteome was characterized (Figure 4B, S4B). First, comparison of *gfp*-GLKD transcriptomes between non-GSCLC and GSCLC contexts revealed that TE transcript levels are generally elevated in GSCLCs (Figure S4C). This baseline increase may account for the relatively modest TE derepression observed for *aub*– or *piwi*-GLKD in GSCLC setting (Figure 4AB).

**Figure 4.**
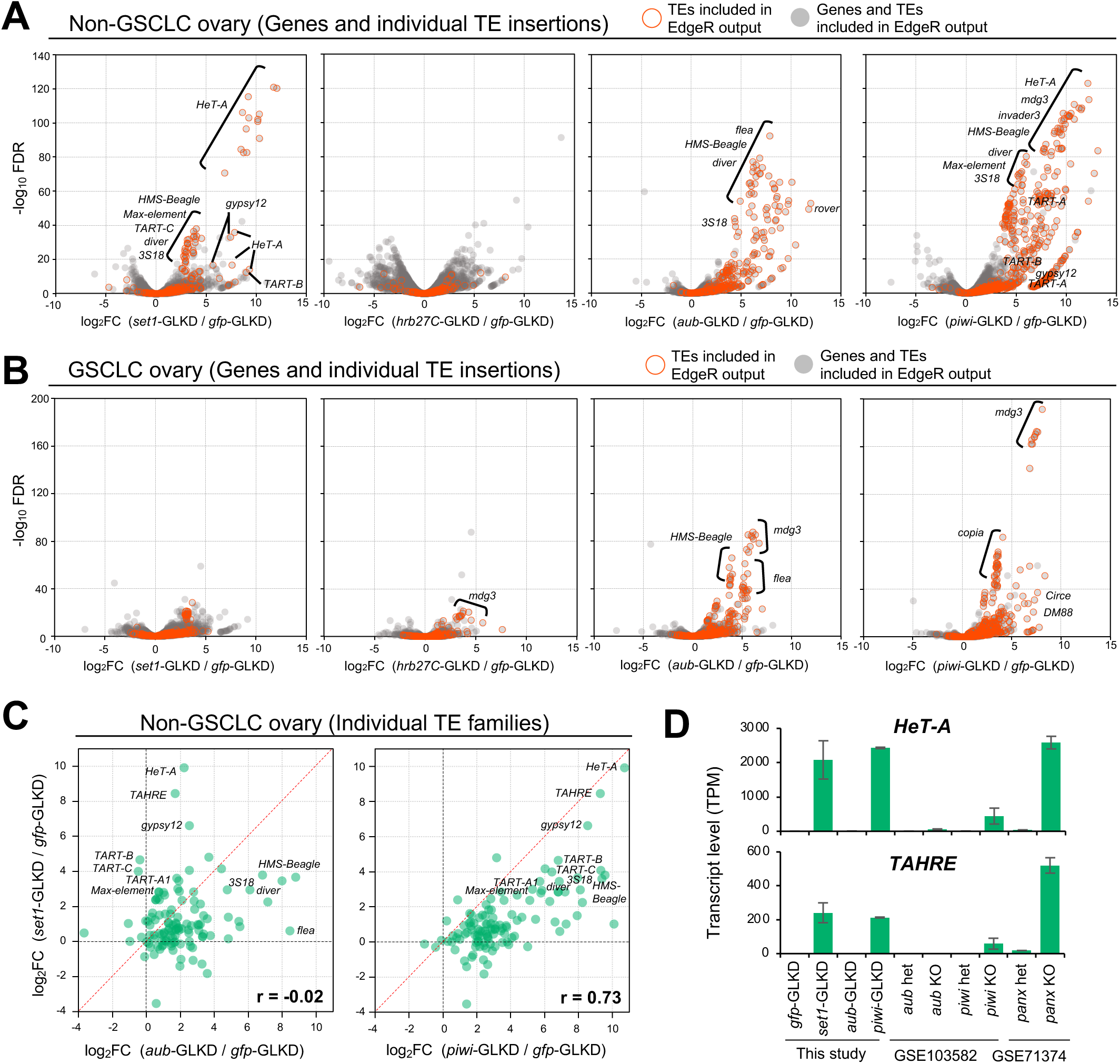
Germline knockdown (GLKD) effect on ovary transcriptome. (A-B) Transcriptome analysis and comparison in non-GSCLC ovaries (A) or in GSCLC ovaries (B). Transcript levels (TPM) of genes and TE insertions in *set1*-, *hrb27C*-, *aub*-, or *piwi*-GLKD condition were compared with those in *gfp*-GLKD control. Dots in the volcano plots; genes (gray) and TEs (gray with red outline) included in the differential expression analysis (EdgeR). (C) Scatter plot comparing the effect of GLKD on the transcript levels of individual TE families (consensus sequence mapping). Left: *set1*-GLKD compared with *aub*-GLKD. Right: *set1*-GLKD compared with *piwi*-GLKD. Fold change from control *gfp*-GLKD (TPM/TPM) were shown. r = pearson correlation coefficient. (D) Transcript levels (TPM) for *HeT-A* and *TAHRE* in different transcriptome datasets. GSE103582 including *aub* and *piwi* knockouts, and GSE71374 including *panx* knockout were analyzed together(Yu *et al*, 2015a; Teixeira *et al*, 2017b). Error bars indicate ±standard deviation (SD) of biological duplicate data.

Hrb27C was identified as the proximity factor of Aub (Figure 2). However, unlike *aub*-GLKD, *hrb27C*-GLKD did not result in clear TE derepression in non-GSCLC ovaries (Figure 4A, S4A). In contrast, TE derepression was evident in GSCLC ovaries (Figure 4B, S4B), and the patterns were positively correlated between *hrb27C-* and *aub*-GLKD conditions (r = 0.57, Figure S4D). These suggest a functional link between Hrb27C and Aub in GSCs.

For Set1 identified as the proximity factor of Piwi (Figure 2), its GLKD caused remarkable derepression of a subset of TEs in non-GSCLC ovaries (Figure 4A, S4A). These *set1*-sensitive TEs included *HeT-A*, consistent with its Gag protein accumulation (Figure 3), as well as *TART* family members, which together with *HeT-A* maintain telomeres as *HTT* array (Villasante *et al*, 2007). *TAHRE*, another *HTT* component, was not analyzed due to the lack of annotated insertions. In addition to *HTT*, several LTR retrotransposons including *HMS-Beagle, Max-element*, *diver*, *3S18*, and *gypsy12* exhibited robust derepression (Figure 4A). This result was corroborated by analyses based on mapping reads to consensus sequences of individual TE families (Figure 4C). Overall, the derepression profile of *set1*-GLKD showed a strong positive correlation with that of *piwi*-GLKD (r = 0.73), but not with *aub-*GLKD (r = 0.02, Figure 4C). Consistent with this, re-analysis of publicly available dataset (GSE103582) indicated that *HTT* members are more sensitive to the loss of *piwi* than that of *aub* (Figure 4D, S4E) (Teixeira *et al*, 2017a). Moreover, similar to *set1*-GLKD, *HTT* members were strongly derepressed upon loss of Panoramix (Panx), an essential cofactor of Piwi (GSE71374) (Sienski *et al*, 2015b; Yu *et al*, 2015a). In contrast to the pronounced effect observed in germline, knockdown of *set1* in ovarian follicle cells using *traffic jam-*Gal4 did not derepress *zam* and *gypsy*, which are silenced by *piwi* in this somatic lineage (Figure S4F). Taken together, these results suggest that Set1 function is associated with Piwi-mediated TE silencing in germline cells.

### Catalytic activity-independent role of Set1 in TE silencing

The C-terminal SET domain of Set1 is responsible for H3K4me3 modification (Mohan *et al*, 2011; Ardehali *et al*, 2011; Wilson *et al*, 2002). To investigate whether this methyltransferase activity is required for TE silencing, we performed rescue experiments by expressing RNAi-resistant transgene encoding GFP-tagged wild-type Set1 (Set1^WT^) or a catalytically inactive variant (Set1^E1613K^) in germline cells depleted of endogenous Set1 (*set1*-GLKD) (Figure 5A) (Vidaurre *et al*, 2024; Hallson *et al*, 2012). Both GFP-Set1^WT^ and GFP-Set1^E1613K^ proteins accumulated in the germline nuclei (Figure 5B). Weaker GFP signals observed for E1613K variant suggest reduced protein stability.

**Figure 5.**
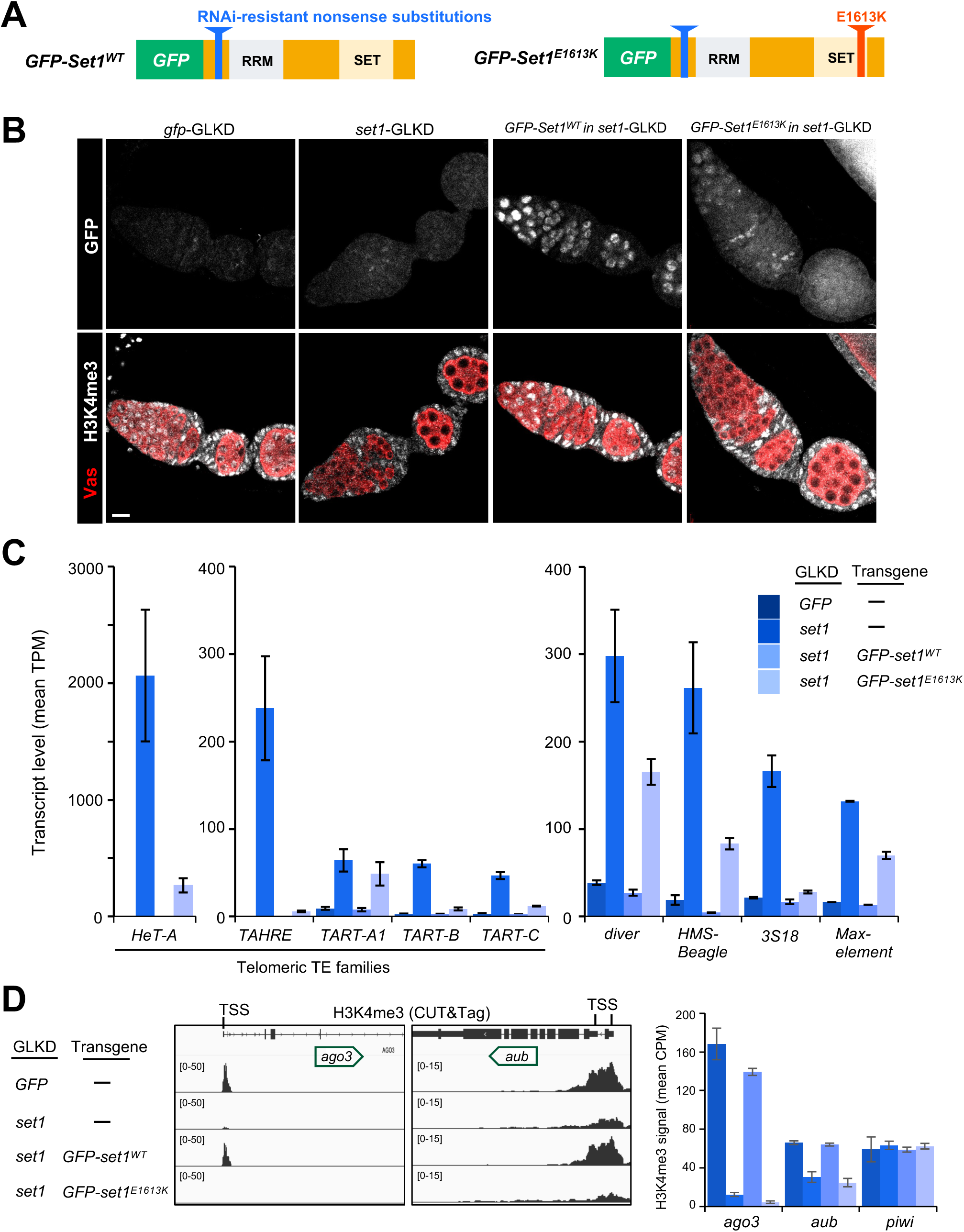
Effect of *set1* transgene expression on H3K4me3 modification and TE expression in *set1*-GLKD ovaries. (A) Schemetic of GFP-Set1 transgenes (Vidaurre *et al*, 2024). Nucleotide substitution to replace 1613^th^ glutamate (E) of Set1 polypeptide to lysine (K) abolishes catalytic activity. Nonsense substitutions render resistance to siRNA targeting. Both placed downstream of UASp and expressed by germline driver NN. (B) GFP fluorescence (top) and immunofluorescence staining for H3K4me3 and Vas (bottom) in egg chambers. Vas marks germline cells. Scale bar = 10μm. (C) Transcript levels (TPM) of TE families in *gfp*-GLKD, *set1*-GLKD, and transgene expression conditions (*GFP-Set1^WT^* or *GFP-Set1^E1613K^* expressed in *set1*-GLKD). Error bars indicate ±SD from biological duplicates. (D) CUT&Tag analysis of H3K4me3 in ovaries of indicated conditions. Left: track view at the *ago3* and *aub* gene loci. Bar graph on the right shows H3K4me3 signals (mean CPM) around TSS (±500bp) of *ago3*, *aub* and *piwi*. Error bars indicate ±SD from biological duplicates.

Transcriptome analysis revealed that *HTT* and other TE families were fully re-repressed by GFP-Set1^WT^ introduced in the *set1*-GLKD background, indicating the depletion of *set1* is causal for their derepression (Figure 5C). Remarkably, although less effective than GFP-Set1^WT^, expression of GFP-Set1^E1613K^ led to substantial re-repression of TEs. This effect was most pronounced for *HTT*, with *TART-A1* as an exception (Figure 5C). Together, these results suggest that Set1 plays a noncanonical, catalytic activity-independent role in TE silencing.

To further support the above notion, we analyzed H3K4me3 signals in *set1*-GLKD and the transgene-rescue conditions. Ovary immunotaining showed that H3K4me3 signals were depleted from germline nuclei in *set1*-GLKD (Figure 5B). Germline H3K4me3 signals were fully recovered by GFP-Set1^WT^, while no recovery was observed with GFP-Set1^E1613K^. We further employed CUT&Tag to genome-widely characterize H3K4me3 signals. H3K4me3 peaks were identified around the transcription start site (TSS) of genes including *ago3* and *aub* (Figure 5D). Consistent with the immunostaining data, H3K4me3 signals on *ago3* and *aub* were markedly reduced in *set1*-GLKD ovaries, and fully restored by GFP-Set1^WT^, but not by GFP-Set1^E1613K^ (Figure 5D). These results confirm the catalytic inactivity of E1613K variant and further support the catalytic activity-independent role in TE silencing. Notably, despite these changes in H3K4me3 signals, steady-state transcript levels of piRNA pathway components were only modestly affected in *set1*-GLKD ovaries (Figure S5A). Moreover, RT-qPCR analysis showed that depletion of COMPASS subunits led to milder derepression of *HeT*-A compared with that of *set1* (Figure S5B). Taken together, these results argue against the notion that Set1 functions solely as an H3K4me3 writer and represses TEs simply by promoting the expression of piRNA pathway components.

### Loss of TE-targeting piRNAs in *set1-*depleted ovaries

TE derepression pattern suggests the functional link between Set1 and Piwi (Figure 4, S4). Moreover, consistent with their close proximity (Figure 2), Set1 and Piwi interact physically, as Piwi co-precipitated with GFP-Set1^WT^ and GFP-Set1^E1613K^ expressed in germline cells (Figure 6A). Notably, the interaction appears transient or weak, given the dependency on crosslinking (Figure S6A). These physical and functional links prompted us to examine whether piRNA expression is affected by *set1* depletion. Deep-sequencing of ovarian small RNAs showed that overall abundance of piRNAs (23∼29-nt fragments and those derived from TE sequences) was largely unchanged in *set1*-GLKD (Figure 6B). However, analysis at the level of individual TE families revealed a striking loss of piRNAs mapping to telomeric *HTT* members (Figure 6C). Notably, antisense piRNAs, which directly target cognate TE transcripts, were reduced more severely than sense piRNAs. A prominent example is *HeT-A*, for which sense piRNAs were largely unaffected whereas antisense species were nearly lost (Figure 6CD). Supporting the link to Piwi pathway, re-analysis of published small RNA-seq data (GSE71374) revealed a similar antisense-biased reduction of *HTT*-mapping piRNAs in ovaries lacking Panx, a key cofactor of Piwi (Figure 6E) (Yu *et al*, 2015a; Sienski *et al*, 2015b). An antisense-biased reduction of piRNAs was also observed for other Set1-controlled TEs, including *3S18*, *gypsy12, Max-element*, *diver*, and *HMS-Beagle* (Figure 6C). Despite this loss, the ping-pong signature (10-nt overlap frequency) of piRNAs was preserved (Figure S6). This is consistent with the maintenance of nuage structure in *set1*-GLKD ovaries (Figure 3B). Together, these findings indicate that Set1 is required to express TE-targeting functional piRNAs, supporting its crucial role in Piwi-mediated TE silencing.

**Figure 6.**
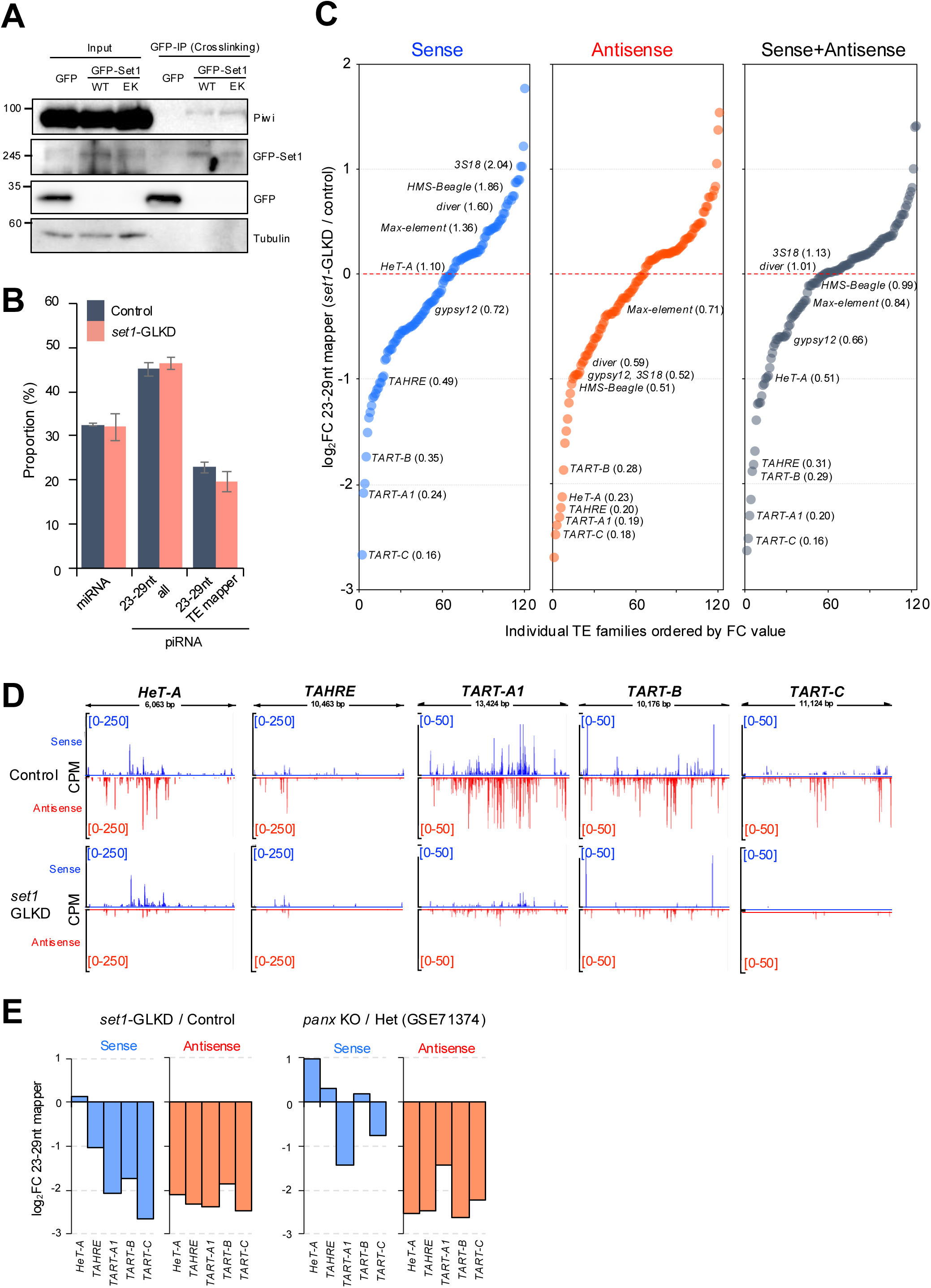
Co-precipitation of Piwi with Set1, and effect of *set1*-GLKD on piRNA accumulation in ovaries. (A) Immunoprecipitation of GFP or GFP-Set1 proteins from ovaries using anti-GFP antibody (GFP-IP) after crosslinking treatment. Tubulin serves as loading control. (B) Proportion (%) of miRNA and piRNA in ovarian small RNAs. miRNA: reads mapping to miRNA precursors. piRNA: 23∼29-nt reads (excluding rRNA, tRNA, snoRNA, snRNA, and miRNA mappers), or 23∼29-nt reads mapping to TE consensus sequences. Control: mean values given by single replicate of *gfp*-GLKD and *GFP-Set1^WT^*-rescue conditions. *set1*-GLKD: mean values given by biological duplicate of *set1*-GLKD condition. Error bars indicate ±SD of biological duplicates. (C) Fold changes (FC) of piRNA levels (RPM) in *set1*-GLKD compared to control condition. piRNA: 23∼29-nt mappers on individual TE families. TEs having few mappers (RPM <5) are excluded. Sense or antisense piRNA: mappers having TE sense strand or the reverse complementary sequences. (D) Track view for 23∼29-nt mappers on consensus sequences of telomeric TE families. (E) Fold change (FC) of the levels (RPM) of piRNA mapping to telomeric TE families. Small RNA libraries for *panx* KO and the heterozygous control conditions (GSE71374) are included (Yu *et al*, 2015a).

### Set1 binds TE sequences and localizes to subtelomeric piRNA cluster loci

How does Set1 contribute to the expression of antisense piRNAs? To address this question, we analyzed the chromatin binding by GFP-Set1 expressed in *set1*-depleted germline cells, using CUT&Tag with anti-GFP antibodies. In parallel with GFP-Set1^WT^, GFP-Set1^E1613K^ was included in the analysis, given the functionality in TE silencing (Figure 5). Comparison between GFP-Set1 and GFP control conditions identified Set1-binding peaks (p<0.01, Figure S7A). Notably, genes and TEs associated with peaks from Set1^WT^ or Set1^E1613K^ showed limited overlap (Figure 7A). However, this does not indicate their targets are distinct; rather, it reflects differential affinity between Set1^WT^ and Set1^E1613K^ to shared targets (r=0.85, Figure 7B). Consistent with its role as H3K4me3 writer, targets identified from Set1^WT^ peaks were predominantly endogenous genes (541 of 548, Figure 7A). These bindnig peaks appeared around transcription start site (TSS) and coincided with *set1*-dependent H3K4me3 signals, as exemplified with *aub* (Figure 7C). In contrast, targets identified from Set1^E1613K^ peaks were enriched with TEs (Figure 7A, 84 of 306), including *HeT-A, TART*, *diver*, *HMS-Beagle*, and *Max-element*, regulated by Set1 (Figure 4A, 5C). At the level of consensus sequences, two major peaks were typically observed: one spanning the 5’UTR-ORF1 region and another within the 3’UTR (Figure 7D, S7CD). Collectively, these results revealed the affinity of Set1 to TE sequences.

**Figure 7.**
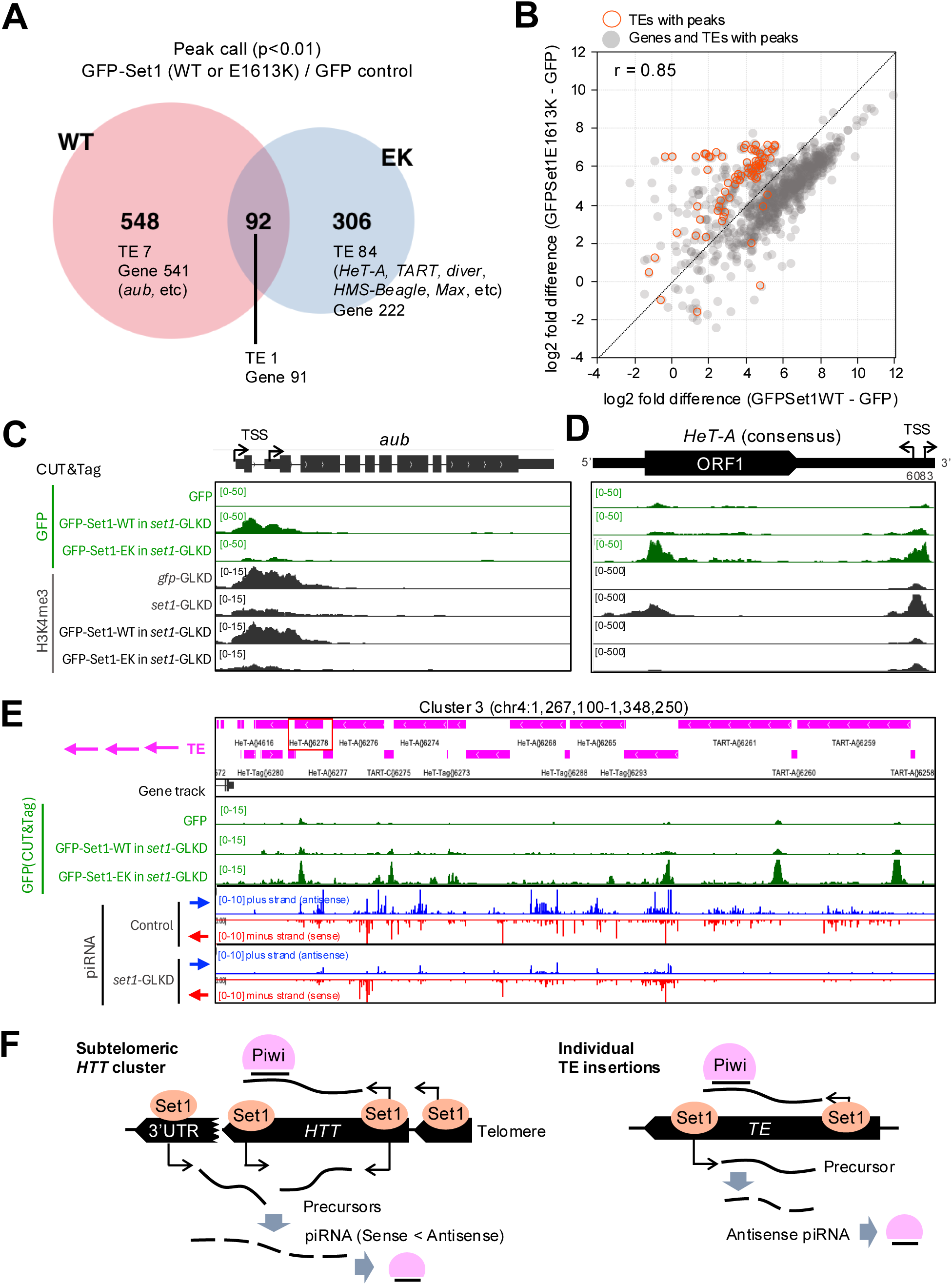
Genome-wide Set1-binding analysis, and a model for Set1 function in TE silencing. (A) Venn diagram for genes and TEs having GFP-Set1^WT^v or GFP-Set1^E1613K^-binding peaks. Peak calling by MACS2 (p<0.01). (B) Scatter plot comparing fold differences of CUT&Tag signals (GFP-Set1^E1613K^ – GFP GFP-Set1^WT^ – GFP) on genes and TEs identified by peak calling. r = pearson correlation coefficient. (C, D) Track view for CUT&Tag signals of GFP proteins (green) and H3K4me3 (black). *aub* (C) and *HeT-A* (D) represent genes or TEs, respectively. TSS: transcription starting site. (E) Track view for CUT&Tag signals of GFP proteins (green) and piRNA levels on subtelomeric piRNA cluster 3. piRNAs having genomic plus strand sequence (Blue) or minus strand sequences (Red) are separately shown. (F) A model for Set1-mediated production of antisense-biased piRNAs on subtelomeric cluster loci and on individual TE insertions outside of cluster loci. Subtelomere regions maintain fragments of 3’UTR derived from *HTT*, which are bound by Set1 to initiate antisense transcription. Catalytic activity of Set1 is dispensable. Similar mechanism is applied for antisense piRNA production from potentially mobile TE insertions outside of piRNA cluster loci. Antisense piRNAs are loaded onto Piwi to reinforce transcriptional silencing.

We mention that H3K4me3 signals on TEs behaved differently from those at gene TSS: they increased upon *set1* depletion and decreased upon expression of both Set1^WT^ and Set1^E1613K^, indicating a lack of correlation with Set1’s catalytic activity (Figure 7D, S7CD). Instead, TE-associated H3K4me3 signals were positively correlated with the transcript abundance (Figure 5), suggesting that these marks are deposited either by residual Set1 in the *set1*-depleted germline cells or through Set1-independent mechanisms (Ardehali *et al*, 2011).

3’UTRs of *HTT* members possess promoter and TSS for bidirectional transctiption (Figure 7D) (Danilevskaya & Arkhipova, 1997; Radion *et al*, 2017; Maxwell *et al*, 2006). Intriguingly, *HTT* 3’UTRs interacting with Set1 are maintained as truncated insertions in subtelomeric piRNA cluster loci. *HeT-A{}6278* in cluster 3 is one of those 3’UTR fragments. Alignment to the consensus sequence suggests that *HeT-A{}6278* maintains promoter and antisense TSS but not sense TSS (Figure S7E). piRNA hot spot starting from this 3’UTR fragment implies the involvement in piRNA precursor transcription (Figure 7E). Set1 showed affinity to other sites on cluster 3, which is contrastive to non-telomeric clusters lacking, if not any, Set1-binding signals (Figure S7F). Insertions from *set1*-sensitive LTR retrotransposons are rare within major piRNA clusters. *Max{}2206* is one of exceptions found in *42AB* cluster, but it did not exhibit Set1 binding (Figure S7F). Hence, Set1 binding to cluster loci is biased toward subtelomeric regions, whose piRNA production is extremely *set1*-dependent (Figure 7E). Set1 binding to LTR retrotransposons may instead reflect its affinity for potentially mobile copies located outside piRNA clusters (Wang *et al*, 2018). Collectively, these results support a model in which Set1 binding to TE sequences promotes precursor transcription for piRNA biogenesis in germline cells (Figure 7F).

## Discussion

To deepen the understanding of piRNA-directed TE silencing, this study explored the proximity factors of individual PIWI proteins expressed in female germline cells. The proteome provided additional clues supporting the known interactions, and, more importantly, revealed otherwise unrecognized links between factors in piRNA biogenesis. First, we point out that Zucchini (Zuc), an endonuclease on mitochondrial outer membrane, was identified in the proximity of both Aub and Piwi. These three proteins are together involved in phased piRNA production (Ge *et al*, 2019), but their physical links have not been reported in ovaries. It should be noted, a study conducting TurboID using Zuc as bait did not find Aub or Piwi in its proximity (Nguyen *et al*, 2023). This might be due to the difference of drivers (matalfa-Gal4 vs nos-Gal4), stages of germline cells (differentiating cells vs stem-cell like cells), or bait proteins (Zuc vs PIWI proteins). In addition to Zuc, our data also showed that Nxf3 and Bootlegger (Boot) mediating the export of piRNA precursors are in the proximity of Ago3, but not of Aub (ElMaghraby *et al*, 2019; Kneuss *et al*, 2019). The selective proximity supports the model whereby precursor transcripts delivered by Nxf3 and Boot from nucleus to nuage in cytoplasm are targeted by Ago3-piRNA complexes (Wang *et al*, 2023b).

Hrb27C emerged as a functionally relevant partner of Aub (Figure 2-4). As a member of hnRNP family, Hrb27C regulates the transport, localization, and translation of interacting mRNAs during oogenesis (Yano *et al*, 2004; Goodrich *et al*, 2004; Huynh *et al*, 2004). Both Hrb27C and Aub are intrinsic factors required for GSC maintenance (Ma *et al*, 2017; Rojas-Ríos *et al*, 2017; Rojas-Ríos *et al*, 2024; Finger *et al*, 2023). In this study, we provided evidence supporting the involvement of Hrb27C in Aub-mediated TE silencing in GSCLCs (GSC-like cells induced by depletion of differentiation factor, *bam*) (Figure 4). The cooperation between Hrb27C and Aub in GSCs warrants further detailed investigation.

H3K4 methyltransferase Set1 was identified and characterized as a key factor in Piwi-mediated TE silencing in female germline (Figure 2-7). We initially assumed that, despite the proximity to Piwi, TE silencing by Set1 could be attributable to its general coactivator function (Ardehali *et al*, 2011; Mohan *et al*, 2011). Set1 acting as H3K4me3 writer inside COMPASS potentially underlies the expression of germline genes including piRNA pathway components. However, our results argue against this notion but instead shed light on its specialized role in Piwi pathway. First, *set1* and *piwi* but not *aub* showed similar TE derepression pattern upon depletion from germline cells (Figure 4). Second, *set1* depletion caused severe loss of piRNAs derived from *HTT*, which we found ocurring in *panx*-deficient ovaries disrupting Piwi pathway (Figure 6). Moreover, we demonstrated that Set1 can control TEs, *HTT* members in particular, without its catalytic activity (Figure 5). Last, CUT&Tag revealed the affinity of Set1 to TE sequences, which is associated with piRNA production (Figure 7).

How Set1 recognizes TE sequences has remained elusive. Set1 interacts with phosphorylated C-terminal domain (CTD) of RNA polII in elongation phase, which allows targeted recruitment and H3K4me3 deposition on actively transcribed genes (Ng *et al*, 2003). In our CUT&Tag analysis, H3K4me3 signals within TE sequences were largely overlapped with Set1-binding regions. Although our results did not support a role for Set1 in depositing these TE-associated H3K4me3 (Figure 7D), RNA polII on transcriptionally active TEs could contribute to Set1 recruitment. However, *HeT-A* showed only mild derepression by downregulation of a COMPASS component, Wdr82 that is the orthologue of yeast Swd2 linking Set1 and RNA polII (Figure S5A) (Bae *et al*, 2020). Hence, RNA polII-dependent recruitment of Set1 as part of COMPASS may be insufficient, and additional mechanisms may contribute. The elevated TE binding observed for the catalytically inactive Set1 variant, compared with the wild-type protein, support a possible alternative mode of recruitment (Figure 7B). Given the physical and function links, involvement of Piwi machinery in recruiting Set1 to TEs is a plausible possibility. Our proximity proteome did not identify SetDB1/Eggless, the H3K9me3 writer responsible for heterochromatin formation at Piwi-piRNA target loci (Rangan *et al*, 2011; Osumi *et al*, 2019; Akkouche *et al*, 2017). This absence suggests that our dataset may be biased toward a specific stage at which Piwi cooperates with Set1. Further characterization of additional factors identified by our Piwi proxiome should provide deeper insight into the mode of actions of Set1 in TE silencing.

Most piRNAs are derived from defined genomic regions called piRNA cluster loci (Brennecke *et al*, 2007b). The heterochromatic structure of these loci requires noncanonical RNA polII recruitment for piRNA precursor transcription, mediated by H3K9me3 reader Rhino and the cofactors (Klattenhoff *et al*, 2009; Mohn *et al*, 2014; Andersen *et al*, 2017). Recent studies uncovered cluster-specific molecular underpinnings for piRNA precursor transcription. These include DNA-binding protein, Kipferl, and enhancer of zeste, E(z), the writer of H3K27me3, each required at distinct cluster loci (Akkouche *et al*, 2025; Baumgartner *et al*, 2022). In gonadal soma, Traffic jam (Tj), a fly orthologue of large Maf transcription factors, activates soma-restricted cluster, *flamenco* (Rivera *et al*, 2025; Alizada *et al*, 2025). Subtelomeric piRNA cluster loci involve Set1 (this study) and NSL complex (Iyer *et al*, 2023). Together, these findings indicate that precursor transcription and piRNA production are governed by diverse chromatin modifiers and transcriptional regulators. Such diversity may be advantageous, enabling spaciotemporal control of the piRNA repertoire to support multitasks of a common set of PIWI proteins. By localizing to subtelomeric cluster loci, Set1 would exert its role in telomere control with Piwi. Characterization of telomere phenotypes in *set1*-depleted germline cells will be an important direction for future studies. Beyond piRNA clusters, Set1 also exhibited affinity for several TE families including *HMS-Beagle* and *Max-element*, both of which maintains mobility in female germline and transmits new copies to next generations through oocyte targeting (Yang *et al*, 2023; Wang *et al*, 2018). The role of Set1 in silencing of such active TEs also represent a future avenue.

Fly Set1 can silence TEs forming telomeres regardless of its methyltransferase activity (Figure 5). This finding is reminiscent of molecular features reported for yeast Set1, which was originally identified as a transcriptional repressor in the telomere and silent mating loci (Nislow *et al*, 1997). Notably, its repressive functions on genes and TEs are often associated with H3K4 methylation-independent mechanisms (Jezek *et al*, 2023; Lee *et al*, 2018; Lorenz *et al*, 2012). However, the mechanisms underlying TE silencing are different between the two organisms, as yeast lacks PIWI clade proteins and does not produce piRNAs. Nonetheless, Set1-dependent antisense non-coding RNAs have been characterized as repressors of retrotransposons in yeast (Berretta *et al*, 2008). The role of Set1 in generating antisense, silencing-competent transcripts for genome defense may be conserved across broader contexts.

## Materials and Methods

### Fly stocks and culture

All stocks and crosses were raised at 25°C on a molasses/yeast medium [5% (w/v) dry yeast, 5% (w/v) corn flower, 2% (w/v) rice bran, 10% (w/v) glucose, 0.7% (w/v) agar, 0.2% (v/v) propionic acid, and 0.05% (v/v) p-hydroxy butyl benzoic acid]. Fly stocks used in this study are listed in **Table S4**.

### Plasmid construction and generation of transgenic flies

All the primers used for plasmid construction are listed in **Table S5**. To generate UASp-mTurbo-GFP, UASp-mTurbo-GFP-Piwi, UASp-mTurbo-GFP-Aub and UASp-mTurbo-GFP-Ago3, mTurbo fragment was amplified by PCR using attB_miniTurbo_Fw_general and miniTurbo_linker_Rv_general as primers and 3xHA-miniTurboNLS_pCDNA3 (addgene #107172) as template. GFP fragment was amplified by PCR using 2-1vLinker>GFP5_F and Ti876-GFP-G5-R as primers. Piwi, Aub or Ago3 fragment was amplified by PCR using primer pairs of Linker_to_Piwi and Piwi_to_Vector, Ti893-G5-Aub-CDS-F and Ti894-Aub-UASp-R1, or Linker_to_Ago3 and Ago3_to_Vector, respectively, with cDNA from *yw* ovaries as template. PCR fragments were introduced into the XbaI site in pUASp-K10-attB vector(Koch *et al*, 2009) using In-Fusion HD Cloning (Takara). UASp-mTurbo-GFP and UASp-mTurbo-GFP-Ago3 were integrated to attP40, while UASp-mTurbo-GFP-Piwi and UASp-mTurbo-GFP-Aub were integrated to attP2, and attP-3B sites, respectively.

### Proximity-dependent biotin pulldown in ovarian germline cells

After eclosion, females expressing mini(m)Turbo-GFP-PIWI proteins were reared at 25°C for 2-3 (non-GSCLC ovaries) or 5-7 days (GSCLC ovaries) in the modified molasses/yeast medium supplemented with 100 μM biotin (Nacalai). For individual conditions, ∼300 ovaries from 150 progenies were collected. Ovaries were homogenized in 150 μL of PI-lysis buffer [50 mM tris-HCl (pH 7.5), 500 mM NaCl, 2 mM EDTA, 2 mM dithiothreitol (DTT), 0.4% (w/v) SDS, and cOmplete Protease Inhibitor Cocktail Tablet (Roche)], using Bioruptor (Diagenode) for 30s (power H) for six times with 30s intervals. Triton X-100 was then added to samples at a final concentration of 2% (v/v), and homogenization was further performed for 30s for three times with 30s intervals. After centrifugation (20,000*g*, 10 min, 4°C), the supernatant was diluted with equal amount of 50 mM Tris-HCl (pH 7.5) buffer and incubated with pre-equilibrated 15-μL slurry volume of Dynabeads MyOne Streptavidin C1 (Thermo Fisher Scientific), overnight at 4°C with gentle rotation. Next day, beads were washed twice with W1 buffer [50 mM tris-HCl (pH 7.5), 250 mM NaCl, 0.2% (w/v) SDS, 1 mM EDTA, and 1 mM DTT], twice with W2 buffer [50 mM HEPES-KOH (pH 7.4), 500 mM NaCl, 0.1% (w/v) deoxycholate (Nacalai), 1% (v/v) Triton X-100, and 1 mM EDTA], twice with W3 buffer [10 mM tris-HCl (pH 8.0), 250 mM LiCl, 0.5% (w/v) deoxycholate, 0.5% (v/v) NP-40 (Nacalai), and 1 mM EDTA], and four times with W4 buffer [50 mM tris-HCl (pH 7.5) and 50 mM NaCl]. The purified proteins bound to beads were stored at –80°C until analysis.

### Immunoblotting and biotinylated protein detection

Protein samples were denatured at 95°C for 5 min in 2× protein loading buffer [4% (w/v) SDS, 200 mM DTT, 0.1% (v/v) bromophenol blue (BPB), and 20% (v/v) glycerol] saturated with biotin, resolved by SDS–polyacrylamide gel electrophoresis (SDS-PAGE) and transferred to 0.2-μm polyvinylidene difluoride membrane (Wako) using the semi-dry system (Trans-blot Turbo, Bio-Rad). The membrane was blocked in 4% (w/v) skim milk (Nacalai) in 1× phosphate-buffered saline (PBS) supplemented with 0.1% (v/v) Tween 20 and further incubated with primary antibodies: rabbit anti-GFP (1:1000, Clonetech), mouse anti-Piwi (1:100) (Saito *et al*, 2006), guinea pig anti-Aub (1:1000) and rat anti-Ago3 (1:200) (Lim *et al*, 2022), mouse anti-Tubulin (1:3000, Santa Cruz). Secondary antibodies were anti-guinea pig immunoglobulins-HRP (1:1000; Dako), anti-rabbit immunoglobulin G (IgG)–HRP (1:3000; Bio-Rad), anti-mouse IgG-HRP (1:3000; Bio-Rad), and anti-rat IgG (1:1000; DAKO). HRP-conjugated Streptavidin (Clonetech) was used for detecting biotinylated Proteins. The chemiluminescent signals generated with Chemi-Lumi One (Nacalai) were detected by Chemidoc MP Imaging system (Bio-Rad). The images were processed with Fiji.

### Proteomic Sample Preparation and Label-Free Quantification using Orbitrap Eclipse Tribrid Mass Spectrometer

Biotinylated proteins bound to magnetic beads were reduced with 1 mM dithiothreitol (DTT) and alkylated with 5.5 mM iodoacetamide. Proteins were digested overnight at 37°C with Trypsin Gold, Mass Spectrometry Grade (Promega, Madison, WI). The resulting peptides were captured and desalted using ZipTip C18 (Millipore, Billerica, MA). Shotgun proteomic analysis was performed using an Orbitrap Eclipse Tribrid mass spectrometer equipped with a FAIMS Pro interface (Thermo Fisher Scientific, Waltham, MA), which was coupled to a Vanquish Neo UHPLC system (Thermo Fisher Scientific, Waltham, MA). Peptides were separated on a reversed-phase column using a linear gradient of 2–24% acetonitrile in 0.1% formic acid at a flow rate of 300 nL/min.

Full MS scans were acquired in the Orbitrap at a resolution of 120,000, followed by MS/MS scans in the ion trap using higher-energy collisional dissociation (HCD) with a normalized collision energy of 35% and a maximum injection time of 10 ms. Label-free quantification (LFQ) was performed using Proteome Discoverer version 2.5 (Thermo Fisher Scientific, Waltham, MA) with the Sequest HT search engine. MS/MS spectra were searched against the UniProt Drosophila melanogaster reference proteome (UP000000803). Protein identifications were filtered at a false discovery rate (FDR) of <1%. Peptide intensities were extracted using the Minora Feature Detector node and used for protein quantification. GO analysis was conducted using MetaScape.(Zhou *et al*, 2019)

### Croslinking and immunoprecipitation

Ovaries were dissected from 50 females expressing GFP, GFP-Set1^WT^, or GFP-Set1^E1613K^ in phosphate-buffered saline (PBS) and immediately fixed with 0.1% (w/v) paraformaldehyde (Electron Microscopy Sciences) for 10 min at room temperature (RT). Fixation was quenched with 125 mM glycine. Samples were then snap-frozen in liquid nitrogen and stored at –80°C until use. Ovary samples were homogenized with a pestle in lysis buffer [20 mM Tris-HCl (pH 7.5), 135 mM NaCl, 1.5 mM MgCl_2_, 10% (v/v) glycerol, 0.2% (v/v) TritonX-100] supplemented with cOmplete Protease Inhibitor Cocktail (Roche). Pre-equilibrated anti-GFP antibody-conjugated magnetic beads (MBL, D153-11) were added to ovarian lysate and incubated for 1h at 4°C. After three times of washing with lysis buffer, bead-bound proteins were extracted by boiling in SDS-PAGE loading buffer [50 mM Tris–HCl (pH 6.8), 2% (w/v) SDS, 100 mM 1,4-dithiothreitol (DTT), 10% (v/v) glycerol, 0.05%(w/v) bromophenol blue].

### Histochemistry and image acquisition

Ovaries were dissected from adult females in 1x PBS buffer supplemented with 0.4% (w/v) bovine serum albumin (BSA; Wako) and fixed in 5.3% (v/v) paraformaldehyde (Nacalai) in 0.67x PBS buffer for 10 min. To observe DNA, ovaries were incubated with 1 μM 40,6-diamidino-2-phenylindole (DAPI) in PBX buffer [1x PBS containing 0.2% (v/v) Triton X-100]. For immunostaining, fixed ovaries were washed with PBX and incubated with Image-iT™ FX Signal Enhancer (Invitrogen) for 30 min and PBX containing 2% (w/v) BSA for 30 min for blocking. The primary antibody incubation was performed overnight at 4°C, and ovaries were washed with PBX at RT for 1h. The secondary antibody incubation was then performed at RT for 2h, and then ovaries were washed with PBX at RT for 1h. The antibodies used for immunostaining were rabbit anti-HeT-A Gag (1:2000),(Lin *et al*, 2023) guinea pig anti-Krimp (1:2000),(Lim & Kai, 2007) mouse anti-Piwi (1:50, Santa Cruz, sc-390946), guinea pig anti-Aub (1:500),(Lim *et al*, 2022) mouse anti-GFP (1:50, Invitrogen, A11120). Secondary antibodies were as follows: Alexa Fluor 488– and 555-conjugated goat antibodies at 1:500 (Molecular Probes) and CF®633 goat antibodies at 1:500 (Biotium). Antibodies were diluted in 0.4% (w/v) BSA containing PBX as the working solution. Images were taken by ZEISS LSM 900 using C-Apochromat 40x/1.20 W Korr objective lens and processed with ZEISS ZEN 3.0 and Fiji.

### Reverse transcription and qPCR analysis

Total RNA was extracted from ovaries using TRIzol™ LS (Invitrogen) following the manufacturer’s instructions. Using DNase I (NEB)–treated RNA, cDNA was synthesized with 2.5 μM oligo(dT) adaptor using SuperScript III reverse transcriptase (Thermo Fisher Scientific). Quantitative reverse transcription PCR (qPCR) reaction was performed using SYBR™ Green qPCR Master Mix (Thermo Fisher Scientific) and gene-specific primers (**Table S5**) in QuantStudio 5 Real-Time PCR system (ABI).

### Transcriptome analysis

Total RNA was extracted from 100 ovaries (non-GSCLC ovaries or GSCLC ovaries) using TRIzol™ LS (Invitrogen). To obtain GSCLC ovaries expressing shRNA for *aub*, *hrb27C*, *set1* or *gfp*, shRNA lines were combined (*bam* shRNA; *aub* shRNA, *bam* shRNA; *hrb27C* shRNA, *bam* shRNA; *set1* shRNA, *bam* shRNA; *gfp* shRNA) and crossed with NN. DNase I-treated RNA was sent to Rhelixa (Japan) for library construction and deep-sequencing. Polyadenylated RNA selection was performed with NEBNext Poly(A) mRNA Magnetic Isolation Module (NEB). Libraries were constructed with the NEBNext Ultra II Directional RNA Library Prep Kit (NEB) and sequenced by using NovaSeq 6000 (Illumina). Trimming was performed by Cutadapt (v1.18, –j 12 –a AGATCGGAAGAGCACACGTCTGAACTCCAGTCA –A AGATCGGAAGAGCGTCGTGTAGGGAAAGAGTGT, –m 20). Trimmed paired-end reads were mapped to the genome (BDGP6.46, dm6) using STAR (v2.7.10b, –-outFilterMultimapNmax 100 –-outSAMmultNmax 1 –-outMultimapperOrder Random). Read counting was performed with featureCounts (-M –p –-countReadPairs –t exon –g gene_id) using Drosophila_melanogaster.BDGP6.46.112.gtf (ensembl.org). Using the sum of RPK (read per kilobase) for individual genes and TEs, TPM (transcript per million) was calculated. Trimmed paired-end reads were also mapped to consensus sequences of individual TE families (transposon_sequence_set.embl.v.9.41, flybase) with STAR using the same options. Mappers on individual TE families were counted using pileup.sh (BBMap). TPM were calculated using the sum of RPK for genes and TEs obtained in genome mapping. Published small RNA data (GSE103582 and GSE71374) were processed in the same way.

### Small RNA sequencing and analysis

Biological duplicate dataset for Set1-present condition was provided by *gfp-GLKD* ovaries (control RNAi condition), and *set1*-GLKD ovaries expressing GFP-Set1^WT^ (rescue condition). Dataset for Set1-depleted condition was provided by *set1*-GLKD ovaries, and the siblings of rescue conditions (CyO). Total RNA was extracted using TRIzol™ LS (Invitrogen) from 60 to 100 ovaries of ∼3 days old adult females. After the addition of chloroform and centrifugation (12,000g 15min, 4°C), short (< ∼200 nt) RNA in the aqueous phase was purified using RNA Clean & Concentrator 5 (R1015 Zymo). To deplete 2S ribosomal RNA (rRNA), 2 pmol of complementary oligo DNA (5’-AGTCTTACAACCCTCAACCATATGTAGTCCAAGCAGCACT-3’) was added per 1 μg of RNA. The mixture was heated (95°C, 2min), gradually cooled down to form DNA/RNA hybrid, and then treated with RNase H (NEB) at 37°C for 30min. RNase H was inactivated by heating at 65°C for 20 min. The 2S rRNA-depleted RNA mixture was loaded onto 8 M urea-polyacrylamide gel (12%) and size-separated in parallel with visible RNA ladder (Dynamarker DM253, BioDynamics Laboratory Inc.) in 0.5x tris-borate EDTA buffer. Gel within the range from 20 to 30nt was excised and passed through a small hole opened by needle (23Gx1, TERUMO) at the bottom of 0.5 mL DNA LoBind tubes (Eppendorf) by centrifugation (15,000g). The fine particles were recovered in 2.0 mL DNA LoBind tubes, and RNAs were eluted overnight at 4°C with gentle rotation in the presence of 300 mM NaOAc (pH 5.2). After removing the particles using cellulose 0.22 µm membrane filter Spin-X (Costar, 8160), RNA was precipitated in the presence of 80% (v/v) ethanol and glycogen (40 μg/mL) (Nacalai) for overnight at –20°C. After centrifugation (20,000g, 20min, 4°C), RNA pellet rinsed twice with 80% (v/v) ethanol was resuspended in RNase-free water. Small RNA libraries were prepared using NEBNext® Multiplex Small RNA Library Prep Set for Illumina (NEB, E7300S) following manufacture’s procedure. After 15 cycles of PCR amplification, the libraries were purified using MagMAX™ Pure Bind Beads (Applied Biosystems), size-separated in 3% (w/v) low melting agarose (HydraGene) in the presence of SYBR gold (Thermo Fisher). ∼150-bp library fragments containing 20∼30-bp inserts were purified using QIAquick gel extraction kit (Qiagen). Libraries were sequenced by RIMD NGS core facility (The University of Osaka) using the NovaSeqX Plus platform (Illumina). Adaptor sequences (AGATCGGAAGAGC) were removed by Trim Galore (v0.6.2, –-length 15) with quality filter cut off (Phred ≥20). Adaptor-trimmed reads were mapped to rRNA, tRNA, snoRNA, and snRNA sequences using bowtie (v1.3.1), and the unmappers were collected (samtools view –f 4). From the unmappers, 18∼29-nt reads were selected as small RNAs using seqkit (v2.4.0) and mapped to miRNA precursors using bowtie (-v 0) to collect “miRNA” populations. The size-selected (23∼29-nt) unmappers were considered as “piRNA”. These piRNA fragments were mapped to TE consensus sequences (transposon_sequence_set.embl.v.9.41, flybase) using bowtie allowing up to 3 mismatches and taking one from multi mappers (-v 3 –M 1 –-best –-strata). Fw and Rv mappers were counted for individual TEs using pileup.sh. Read counts were normalized using the sum of 18∼29-nt reads and reads per million (RPM) was obtained. For visualization, Fw and Rv mappers were separated using splitsam.sh. Bedgraph was generated using bedtools (v2.26.0, genomecov –bga –split), values normalized using scale factors given by the sum of 18∼29-nt reads. Mean values of biological replicates were obtained using bedtools unionbedg. Track view was generated in IGV (v2.16.0). piRNA overlap scores (z) were measured using signature.py (Antoniewski, 2014). piRNA fragments were also mapped to Drosophila genome (BDGP6.46, dm6) using bowtie without allowing mismatch (-v 0 –M 1 –-best –-strata). Bedgraph generation, normalization, averaging, and visualization follow the processes in the analysis using TE consensus sequences. Published small RNA data (GSE71374) were processed in the same way.

### CUT&Tag sequencing and analysis

We referred (Anderson *et al*, 2023) for performing whole ovary CUT&Tag. Twenty ovaries from 10 adult females at ∼3 days old were dissected in PBS buffer supplemented with cOmplete Protease Inhibitor (Roche). Ovaries were permeabilized in PBX buffer (PBX containing 0.2%(v/v) Triton-X100) for 30 min at RT. After removing PBX, ovaries were washed once with Wash+ buffer (20mM HEPES [pH 7.4], 150 mM NaCl, 0.5 mM spermidine[Nacalai], 2 mM EDTA, 1%[w/v] BSA, cOmplete Protease Inhibitor) and incubated overnight at 4°C with primary antibodies diluted by 1:50 with Wash+ buffer. Anti-H3K4me3 (#61979, mouse, Active Motif), or anti-GFP (#598, rabbit, MBL) antibodies were used. Next day, ovaries were washed three times with Wash+ buffer, and incubated with secondary antibodies diluted by 1:50 with Wash+ buffer. Anti-mouse IgG (#52885L, Cell Signalling), or anti-rabbit IgG (#35401S, Cell Signalling) were used. After removing the secondary antibody solution, ovaries were washed three times with 300Wash+ buffer (20 mM HEPES [pH 7.4], 300 mM NaCl, 0.5 mM spermidine, cOmplete Protease Inhibitor) and incubated for 1h at RT with loaded pAG-Tn5 (#79561S, Cell Signalling) diluted by 1:25 with 300Wash+ buffer. After removing the Tn5 solution, ovaries were washed three times with 300Wash+ buffer. Tagmentation was performed by incubating ovaries in 300Wash+ buffer supplemented with 10 mM MgCl_2_ for 1h at 37°C. After removing the supernatant, ovaries were treated with collagenase (2 mg/mL, Sigma C9407) in HEPESCA buffer (50 mM HEPES [pH 7.4], 360 µM CaCl_2_) for 1h at 37°C. Then, the reaction mixture was added with SDS, EDTA, and proteinase K (Nacalai) (final concentration 0.2%[w/v], 16 mM, and 0.3 mg/mL, respectively), and further incubated for 1h at 58°C. After the reaction, DNA was purified by using Quick-DNA Microprep kit (D3020, ZYMO). Using the purified DNA as template, libraries were synthesized by PCR (72°C 5min, 98°C 30s, 14 cycles of [98°C 10s, 63°C 15s], 65°C 5min) using Ultra II Q5 Master Mix (NEB) and dual-indexed primers (Table S5). Generated libraries were purified using x1.3 volume of AmpureXP (beckman). The concentration of libraries was measured using Qubit dsDNA HS assay kit (Thermo). Libraries were sequenced with NovaSeqX (Illumina) and 150+150bp paired-end reads were obtained (H3K4me3 libraries sequenced by RIMD in The University of Osaka, GFP libraries sequenced by Rhelixa). Adaptor trimming was performed by Cutadapt (-a CTGTCTCTTATACACATCTCCGAGCCCACGAGAC –A CTGTCTCTTATACACATCTGACGCTGCCGACGA, –m 20). Trimmed reads were mapped to *D. melanogaster* genome (BDGP6.46, dm6) using Bowtie2 –p 12 –-no-unal –-end-to-end –-very-sensitive –-no-mixed –-no-discordant –q –-phred33 –I 10 –X 700). Read counts were normalized using deeptools (bamCoverage –-normalizeUsing CPM –of bigwig –-binSize=1). Average bigwig was generated from biological duplicate bigwig data using wiggletools and ucsc-wigtobigwig (wigToBigWig). Track view was generated in IGV (v2.16.0). Counting reads on gene TSS±500bp (H3K4me3) or exon (GFP) was performed using featureCounts (v2.1.1, –M –p –-countReadPairs –O) with Drosophila_melanogaster.BDGP6.46.112.gtf (Ensembl). RPM (read per million) was calculated using the number of aligned reads in Bowtie2 mapping. Trimmed reads were also mapped to TE consensus sequences (transposon_sequence_set.embl.v.9.41, flybase). Bigwig was generated using deeptools (bamCoverage), normalizing the values with scale factors given by the sum of aligned reads on the genome. Mean values of biological replicates were obtained using wiggletools and wigToBigWig (ucsc-wigtobigwig). Track view was generated in IGV (v2.16.0).

## Acknowledgement

We thank Dr. Prof. Xin Chen for providing fly stocks. We thank Chisato Yanagisawa, Minako Moriguchi for their helps with maintenance of fly lines. We are also grateful to Bloomington Drosophila Stock Center for providing fly stocks. We also thank all members in our lab for their insightful discussion and suggestions. This study is supported by JSPS Grant-in-Aid for Scientific Research C (22K06081) for TI, Life Science Foundation of Japan (J231503018) for TI, Takeda Foundation (J241503007) for TI, Daiichi Sankyo Foundation of Life Science (J241503010) for TI, Naito Foundation (J241503011) for TI, JSPS Grant-in-Aid for Scientific Research B (21H02401) for TK, JSPS Grant-in-Aid for Transformative Research Areas A (21H05275) for TK, Grant-in-Aid for JSPS Fellows (23KJ1521) for WI, and Grant from Open and Transdisciplinary Research Initiatives (OTRI) RNA Frontier Science Division for TI. The founders had no role in study design, data collection and analysis, decision to publish, or preparation of the manuscript.

## Author contributions

Conceptualization: TI and TK

Methodology: TI, TK, and WI

Investigation: TI, WI, HK-H and MO

Supervisions: TI and TK

Writing-original draft: WI and TI

Writing-review and editing: TI, WI and TK

## Competing interest statement

The authors declare no competing interests.

## Data availability

Newly generated transcriptome data are available with BioProject accession ID: PRJNA1345876; *Drosophila* ovary polyA transcriptome sequencing, PRJNA1345540; *Drosophila* ovary small RNA sequencing, PRJNA1345531; *Drosophila* ovary CUT&Tag sequencing

## Figure legends

**Figure S1.** Optimization of germline PIWI proximity labelling and purification. (A) Comparison of biotinylation efficiency and biotinylated protein purification between non-GSCLC and GSCLC ovaries. Coomassie brilliant blue (CBB) staining serving as protein loading control. Ovaries expressing mTurbo-GFP or mTurbo-GFP-Aub were analyzed with negative control (without mTurbo protein expression). Asterisks: non-specific signals. (B) Schematic illustration for germline differentiation and the function of Bam. Left; the germarium in wild-type, non-GSCLC ovaries. A germline stem cell (GSC, blue) divides to produce a daughter cell (red) which expresses Bam and starts the differentiation. Two-to-three GSCs are normally maintained in a germarium. Right: by *bam*-GLKD, daughter cells cannot differentiate and thus accumulate as GSCLCs. (C) A pair of wild-type ovaries (*y w*, left) and that of *bam*-GLKD ovaries (right). Scale bar = 1 mm. (D) Immunoblotting images comparing the expression level of mTurbo-GFP-tagged Piwi or Aub with that of endogenous Piwi or Aub in GSCLC ovaries.

**Figure S2.** Quality assessment of PIWI proximity proteome data. Volcano plots for the proximity factors of Piwi, Aub, or Ago3 in individual biological replicates. Red dots highlight the factors showing significant enrichment (abundance ratio > 1, adjusted p < 0.05). Statistics relies on background-based t-test.

**Figure S3.** Germline knockdown (GLKD) efficiency and the effect on protein subcellular localization. (A-C) Plotting the fluorescence signals from egg chambers of indicated genotypes, related to Figure 3B. (A) Krimp (green) and DAPI (magenta), (B) Aub (green) and DAPI (magenta), (C) Piwi (green) and DAPI (magenta). (D) Transcript levels of *aub*, *hrb27C* and *set1* in individual GLKD conditions (*ΔΔ*Ct method using *rp49* as reference). Error bars indicate standard deviation (SD) from three biological replicates. *p* values from unpaired Student’s t-test.

**Figure S4.** GLKD effect on ovary transcriptome, and somatic KD effect on TEs. (A-B) Transcriptome analysis and comparison in non-GSCLC ovaries (A) or in GSCLC ovaries (B). Transcript levels (TPM) of genes and TE insertions in *set1*-, *hrb27C*-, *aub*-, or *piwi*-GLKD conditions were compared with those in the control *gfp*-GLKD. Dots in the MA plots; genes (gray) and TEs (gray with red outline) included in the differential expression analysis (EdgeR). (C) Transcript levels (TPM) of individual TE families in *gfp*-GLKD (consensus sequence mapping) compared between non-GSCLC and GSCLC ovaries. (D) Scatter plot comparing the effect of GLKD on the transcript levels of individual TE families (consensus sequence mapping). Left: *hrb27C*-GLKD compared with *aub*-GLKD. Right: *hrb27C*-GLKD compared with *piwi*-GLKD. Fold changes (TPM/TPM) from control *gfp*-GLKD are shown. r = pearson correlation coefficient. (E) Transcript levels (TPM) for *TART-A1*, *TART-B*, and *TART-C*, in different transcriptome datasets. GSE103582 including *aub* and *piwi* knockout conditions, and GSE71374 including *panx* knockout condition were analyzed. Error bars indicate ± SD of biological duplicates. (F) RT-qPCR measurement of TE transcript levels in ovaries upon somatic KD of *gfp*, *piwi* or *set1* using *tj*-Gal4 driver. *ΔΔ*Ct method using *rp49* as reference. Error bars indicate ±SD from three biological replicates.

**Figure S5.** Effect of COMPASS subunit GLKD on *HeT-A* expression, and effect of *set1*-GLKD on the expression of piRNA pathway factors. (A) RT-qPCR measurement of *HeT-A* transcript levels in COMPASS subunit GLKD ovaries. *ΔΔCt* method using *rp49* as a reference. Error bars indicate ±SD from three biological replicates. *p* value from unpaired Student’s t-test is indicated. Asterisk; *p* < 0.05. (B) Bar graph shows the transcript levels (TPM) of piRNA pathway components in *gfp*-GLKD and *set1*-GLKD ovaries. Error bars indicate ± SD of biological duplicates.

**Figure S6.** Immunoprecipitation of GFP-Set1 without crosslinking, and effect of *set1*-GLKD on ping-pong signature of piRNAs. (A) Immunoprecipitation of GFP or GFP-Set1 proteins from ovaries using anti-GFP antibody (GFP-IP) without crosslinking treatment. Tubulin serves as loading control. (B) Ping-pong signature analysis on piRNAs mapping to telomeric TE families. Z10 scores (reflecting 10-nt overlap frequency within piRNA groups) were highlighted. Error bars indicate ±SD of biological duplicates.

**Figure S7.** Genome-wide Set1-binding analysis. (A) Venn diagram shows overlap of genes and TEs between replicate 1 and 2 of GFP-Set1^WT^ CUT&Tag (peak calling using GFP control). (B) Venn diagram shows overlap of genes and TEs between replicate 1 and 2 of GFP-Set1^E1613K^ CUT&Tag (peak calling using GFP control). (C) Track view for CUT&Tag signals of GFP proteins (green) and H3K4me3 (black) on *TAHRE* (left) and *TART-C* (right). (D) Track view for CUT&Tag signals of GFP proteins (green) and H3K4me3 (black) on *HMS-Beagle* (left) and *Max-element* (right). (E) Alignment of *HeT-A{}6278* to *HeT-A* consensus sequence. (F) Track view for CUT&Tag signals of GFP proteins (green) and piRNA levels on *42AB* locus. Blue and Red indicate piRNAs containing genomic plus and minus strand sequences, respectively.

